# A Neurotomographic Approach for Mesoscale Mapping of Neural Circuits

**DOI:** 10.64898/2026.08.11.743991

**Authors:** Oluwamuyiwa A. Ayanshina, Tolulope T. Adeyelu, Michelle L. Osborn, Kenneth L. Matthews, Charles C. Lee

**Affiliations:** Department of Comparative Biomedical Sciences, Louisiana State University, School of Veterinary Medicine, Baton Rouge, LA, USA; Department of Biomedical Sciences, University of Georgia, Athens, GA, USA; Department of Physics and Astronomy, Louisiana State University, College of Science, Baton Rouge, LA, USA

**Keywords:** micro-CT, retrograde tracers, in vivo imaging, colloidal gold, WAHG, mesoscale connections, convergence

## Abstract

**Background:** Brain regions integrate neural information arriving from several convergent projection sources. At the mesoscale level, neural projections can potentially span both hemispheres and extend along the entire rostrocaudal axis, which complicates efforts to map their full extent. To address this issue, we describe a novel method for mapping such mesoscale connectivity *in vivo* and *ex vivo*. Our ‘neurotomographic’ approach utilizes micro-computed tomography (micro-CT) to image the spatial distribution of neural tracers bound to high Z-elements, e.g, gold.

**Methods:** In this study, we conjugated colloidal gold to a retrograde tracer wheat-germ agglutinin apo-horseradish peroxidase (WGA-HRP) and then stereotactically injected the gold-bound tracer (WAHG) into the mouse forebrain. Micro-CT was then used to image the brain *in vivo* and *ex vivo*, followed by three-dimensional reconstruction of tracer distribution. We then validated our approach by histologically processing the brains using silver enhancement to label gold particles; this enabled a direct comparison of histological labeling with the neurotomographic images.

**Results:** We found that micro-CT imaging could reveal the major spatial distributions of the gold-bound tracer, which was consistent across *in vivo* and *ex vivo* imaging conditions. Moreover, the neurotomographically determined patterns corresponded with the labeling observed in histologically processed tissue, with the major sites of labeling reliably detected in reconstructed neurotomographic images.

**Conclusions:** Overall, our findings demonstrate a potential novel method for non-destructive, three-dimensional mapping of neural tracers *in vivo*. This novel approach can potentially guide targeted multi-site recordings, enable validation of injection site placement, and facilitate rapid longitudinal connectomic analyses *in vivo*.

## INTRODUCTION

Classical neuroanatomical tract tracing remains the gold standard for assessing the pattern of mesoscale connections in the brain [1-6]. Conventional tract tracing relies on serially sectioning the injected brains followed by two-dimensional histological processing and subsequent section alignment for volumetric reconstruction [6]. This process is inherently destructive and does not faithfully recapitulate the structure of the brain *in situ* [7]. Histological processing introduces geometric distortions due to sectioning artifacts, tissue shrinkage, warping, and uneven staining [8, 9]. Even with advanced reconstruction protocols, the fidelity of the final image is generally limited by processing deformations that accumulate across multiple sections [10-13]. Moreover, long-range projection pathways often span hundreds of serial slices, which results in the reconstruction process being labor-intensive and error-prone.

These technical issues have been somewhat mitigated with recent whole-volume optical approaches, involving tissue clearing combined with light-sheet microscopy [14-16]. While these approaches mitigate sectioning artifacts by enabling imaging of intact specimens, they are subject to their own drawbacks [17, 18]. Tissue clearing requires extensive chemical processing, which may alter 3D tissue morphology and may be unsuitable for very large samples [14, 15, 17]. Imaging of the cleared tissue then can depend on fluorescence stability and optical transparency of the sample [16, 18]. Moreover, scattering properties can further constrain the resolution in densely labeled or myelinated regions. Thus, despite these methodological advances, there remains a need for a non-destructive imaging strategy that is capable of resolving long-range connectivity in the brain *in vivo* and *ex vivo*.

In this regard, micro–computed tomography (micro-CT) offers a fundamentally different imaging approach, whose contrast mechanism is based on X-ray attenuation rather than optical fluorescence or absorbance [19, 20]. As an imaging approach, micro-CT benefits from an extensive theoretical and methodological framework established over decades in medical physics [19, 21, 22]. The approach enables the 3D structure of the biological specimen to be assessed *in situ*, with the added benefit of digital sectioning in arbitrary orientations without physical slicing. Nevertheless, the approach has not been routinely applied to studies of the brain and spinal cord, due to the low X-ray attenuation of soft tissue [23]. However, such weak tissue attenuation is advantageous for labeling studies, like those used in tract tracing. Specifically, neural tracers can be conjugated with X-ray contrast agents, which are generally high–atomic number (high-Z) elements, such as gold or iodine [24-28]. These elements exhibit strong attenuation coefficients relative to soft tissue and are thus readily detected in micro-CT scans [24].

Here, we describe our approach to visualize globally connected neural circuits in the intact brain *in vivo* and *ex vivo*. We utilized micro-CT to image the whole-brain distribution of gold-coupled retrograde tracers following their injections in the mouse brain. We found that this imaging approach can reliably detect the gold-bound tracer in intact brain specimens, which also corresponded to the major labeling observed in histologically processed specimens. As such, these experiments demonstrate a potential scalable and non-destructive approach for mesoscale circuit reconstruction that complements existing histological and optical methods, with the advantage of identifying connectivity patterns in the living animal. We have termed our approach for imaging neural connections using micro-CT with the portmanteau: *neurotomography*.

## MATERIALS AND METHODS

### Animals

Adult C57BL/6J mice (3-6 mos., both sexes) (Jackson Labs, Bar Harbor, ME) were used in this study. Animals were housed under standard laboratory conditions with *ad libitum* access to food and water. All procedures were conducted in accordance with institutional guidelines and approved by the relevant Institutional Animal Care and Use Committee (IACUC).

### WGA–apoHRP Gold Conjugate (WAHG) Tracer Preparation

A colloidal gold-conjugated to a wheat germ agglutinin tracer (WAHG) was used for micro-computed tomography (µCT)–based neuroanatomical tracing [26-28]. WAHG preparation was derived from the method described by Basbaum and Menterey (1987). In brief, gold chloride was used as the precursor for gold nanoparticle formation [26]. Wheat germ agglutinin conjugated to apo-horseradish peroxidase (WGA–apoHRP) (Sigma, St. Louis, MO) was incorporated as the targeting and enzymatic component. Gold colloids were prepared from gold chloride trihydrate (HAuCl4 *3H2O) in a stabilizing buffer containing sodium citrate, polyethylene glycol (PEG), tannic acid, and sodium azide (Sigma, St. Louis, MO). The reaction mixture was stirred under controlled conditions to promote reduction of gold ions and formation of colloidal gold nanoparticles. WGA–apoHRP was then added to allow adsorption/conjugation onto the gold nanoparticle surface, forming the WGA-apoHRP–gold complex (WAHG) [26]. The reaction mixture was subjected to sequential ultracentrifugation steps to purify and concentrate the WAHG tracer. Samples were centrifuged at 35,000 rpm for 1 hour, and the supernatant was carefully removed. The pellet was resuspended in buffer and centrifuged again at 35,000 rpm for 1 hour to remove unbound protein and excess reagents. A final purification step was performed at 25,000 rpm for 1 hour. The purified tracer (WAHG) was resuspended in sterile buffer and stored at 4°C until use.

### Stereotactic injections

Mice surgeries and injections were performed similar to that previously described [29]. Briefly, mice were anesthetized using ketamine (100 mg/kg)/xylazine (10 mg/kg) at a dosage of 0.1ml/10g, shaved and mounted on a stereotaxic frame, as previously described. Following aseptic technique, a craniotomy was performed over the injection site and 1 µL of WAHG was unilaterally injected into the hippocampus (HPC) (relative to bregma: AP 2.78; ML +3.5; DV -2.2), coordinates determined based on a standard mice brain atlas [30]. Following these injections, the needle was left in place for several minutes before slow withdrawal. The incision was sutured, and animals were allowed to recover under constant monitoring.

### In vivo micro-CT imaging

Seven days following tracer injection, animals underwent *in vivo* micro-CT (µ*CT*) imaging to assess tracer distribution. Images were acquired using Triumph II SPECT-CT, TriFoil Imaging®, USA (Cheyenne, WY). Animals were anesthetized through continuous and controlled exposure to isoflurane during scanning to prevent motion artifacts. Scanning parameters (tube potential 75 kVp, tube current and exposure time 25 µAs, and voxel size 0.05 × 0.05 × 0.05 mm3) were optimized to detect gold-based contrast within brain tissue. Three-dimensional reconstructions were generated from acquired projection data using manufacturer-provided reconstruction software. Regions of tracer accumulation were visualized and analyzed volumetrically, as described below.

### Tissue Perfusion, Post-Fixation and ex vivo µCT Imaging

Following *in vivo* imaging, animals were transcardially perfused with 0.01M phosphate-buffered saline (PBS) followed by a fixative solution, 4% paraformaldehyde in 0.01M PBS azide (Electron Microscopy Sciences, Morgantown, PA). Brains were extracted and post-fixed for an appropriate duration. *Ex vivo* µCT scans were performed on brain specimens *in situ* and *ex situ*, for higher-resolution imaging and for improved signal-to-noise ratio. Imaging parameters were adjusted to maximize detection sensitivity for gold nanoparticle signal. The *ex vivo* micro-CT images were acquired using Scanco-40 µCT (Scanco Medical AG, Wangen-Brüttisellen, Switzerland) (tube potential 55 kVp, tube current and exposure time 44 µAs, and voxel size 0.012 ×0.012 ×0.012 mm^3^).

### Image processing

Micro-CT images were 3-dimensionally reconstructed using the proprietary instrumentation software on the Triumph II SPECT-CT (Cheyenne, WY) or the Scanco-40 µCT (Scanco Medical AG, Wangen-Brüttisellen, Switzerland). The generation of 3D volume rendering, and virtual slices was performed using OsiriX MD (Pixmeo, Bernex, Switzerland) or VivoQuant 2.50 (Invicro, Boston, MA).. Reconstructed volumes were generated using filtered back-projection and aligned to coronal planes following standard neuroanatomical atlases [30].

### Histology and silver enhancement staining

After *ex vivo* imaging, brains were cryoprotected, then sectioned coronally using a cryostat at 50 μm thickness. To enhance visualization of gold nanoparticles at the microscopic level, silver enhancement staining was performed, similar to our prior studies [27, 28, 31]. Silver ions (Ag^+^) were reduced on the surface of WAHG-labeled colloidal gold particles, resulting in amplification of the signal and improved detectability under light microscopy; brain sections were incubated using Aurion R-Gent silver enhancement solution (Aurion R-Gent SE Intense, Netherlands). Stained sections were mounted onto glass slides and imaged by Akoya PhenoImager slide scanner (Akoya Biosciences, Marlborough, MA). Digital images were acquired for qualitative assessment of tracer localization and comparison with µCT findings. Micro-CT and histology images were co-registered using gross anatomical landmarks.

## RESULTS

We first assessed whether micro-CT could reliably detect the gold-bound retrograde tracer (WAHG) *in vivo*. In these experiments, WAHG was injected into the dorsal hippocampus of the mouse brain and the animals were then allowed to recover for 7 days to allow for adequate transport. Following the recovery period, animals were anesthetized and their heads were scanned using an *in vivo* Triumph II micro-CT/SPECT (Fig. 1; Suppl. Video 1). These scans clearly identified the injection site location and distal retrograde labeling; the 3D location of the injection sites corresponded to the targeted coordinates, as estimated by their relative distance from imaged skull sutures (bregma and lambda) (Fig. 1; Suppl. Video 1).

**Figure 1.**
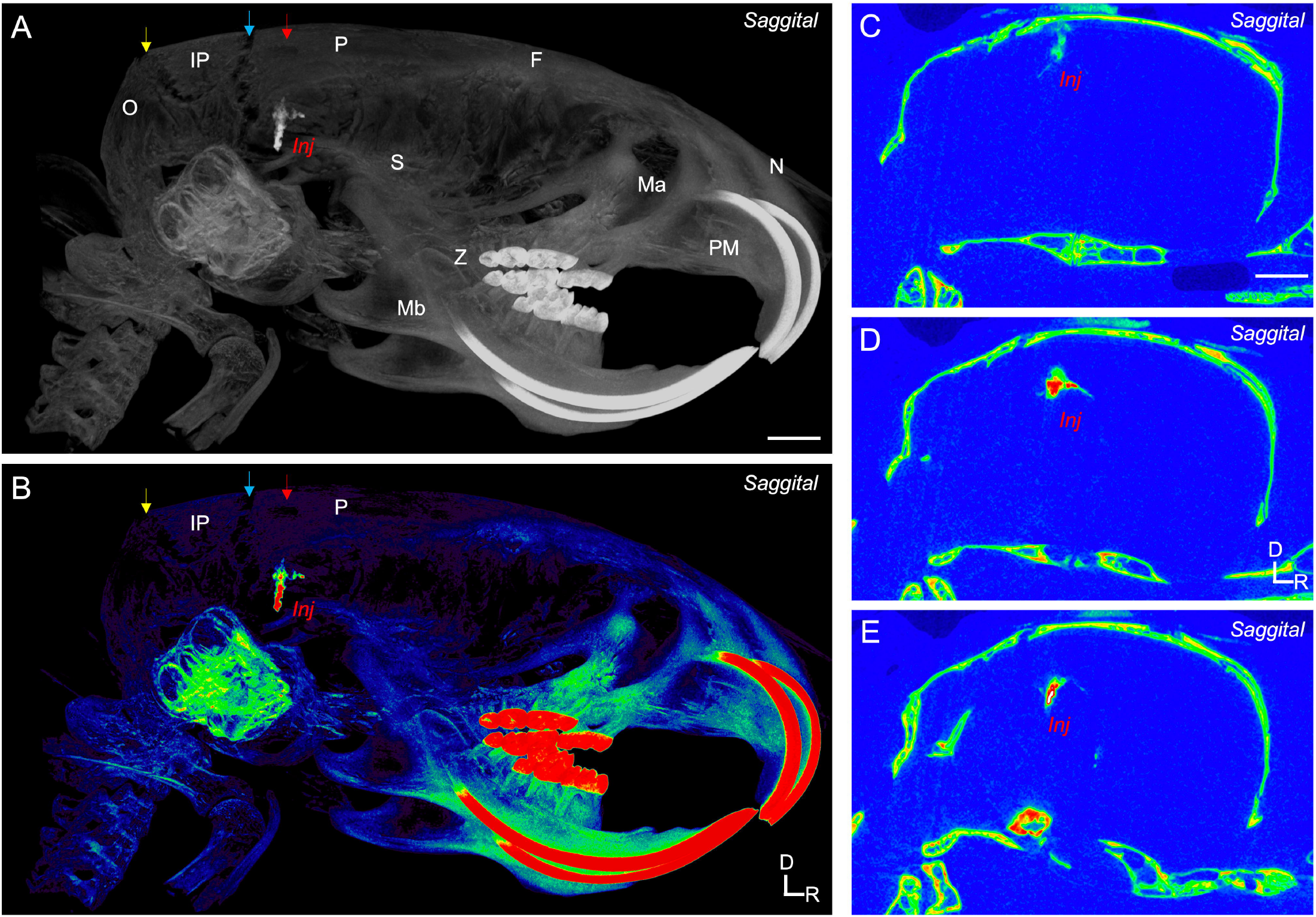
Neurotomographic imaging of gold-bound retrograde tracer (WAHG) injections *in vivo*. **A-B.** 3D-reconstructed micro-CT sagittal view of the mouse skull demonstrating the high intensity absorption of the tracer (*Inj*) within the skull. The retrograde tracer presents with a Hounsfield unit that is coincidentally similar to that of tooth enamel, as shown by similar shades of red in image *B*. Cranial plates indicated by lettered abbreviations. **C-E**. Parasagittal sections demonstrating the distribution of tracer (*Inj*). See also Supplementary Figure 1 and Supplementary Video 1. Scale bars: 1 mm. Orientation: D, dorsal; R, rostral. Red arrows: craniotomy location. Blue and yellow arrows: lambdoid sutures. Cranial plate abbreviations: F, fontal; IP, interparietal; Ma, maxilla; Mb, mandible; N, nasal; O, occipital; P, parietal; PM, pre-maxillary; S, squamosal; Z, zygomatic

We then assessed whether the resolution of acquired images were improved using an *ex vivo* micro-CT scanner, which has a higher beam intensity compared to the *in vivo* scanner (see Methods). Therefore, following the *in vivo* scans, animals were sacrificed to enable the scanning of their heads in the *ex vivo* Scanco µCT 40 desktop cone-beam scanner (Fig. 2i, Suppl. Video 2). As predicted, we observed an increase in the resolution of acquired images in the *ex vivo* scans, with additional fine labeling structure visible surrounding the injection site and in the retrogradely labeled locations (Fig. 2i; Suppl. Video 2).

**Figure 2.**
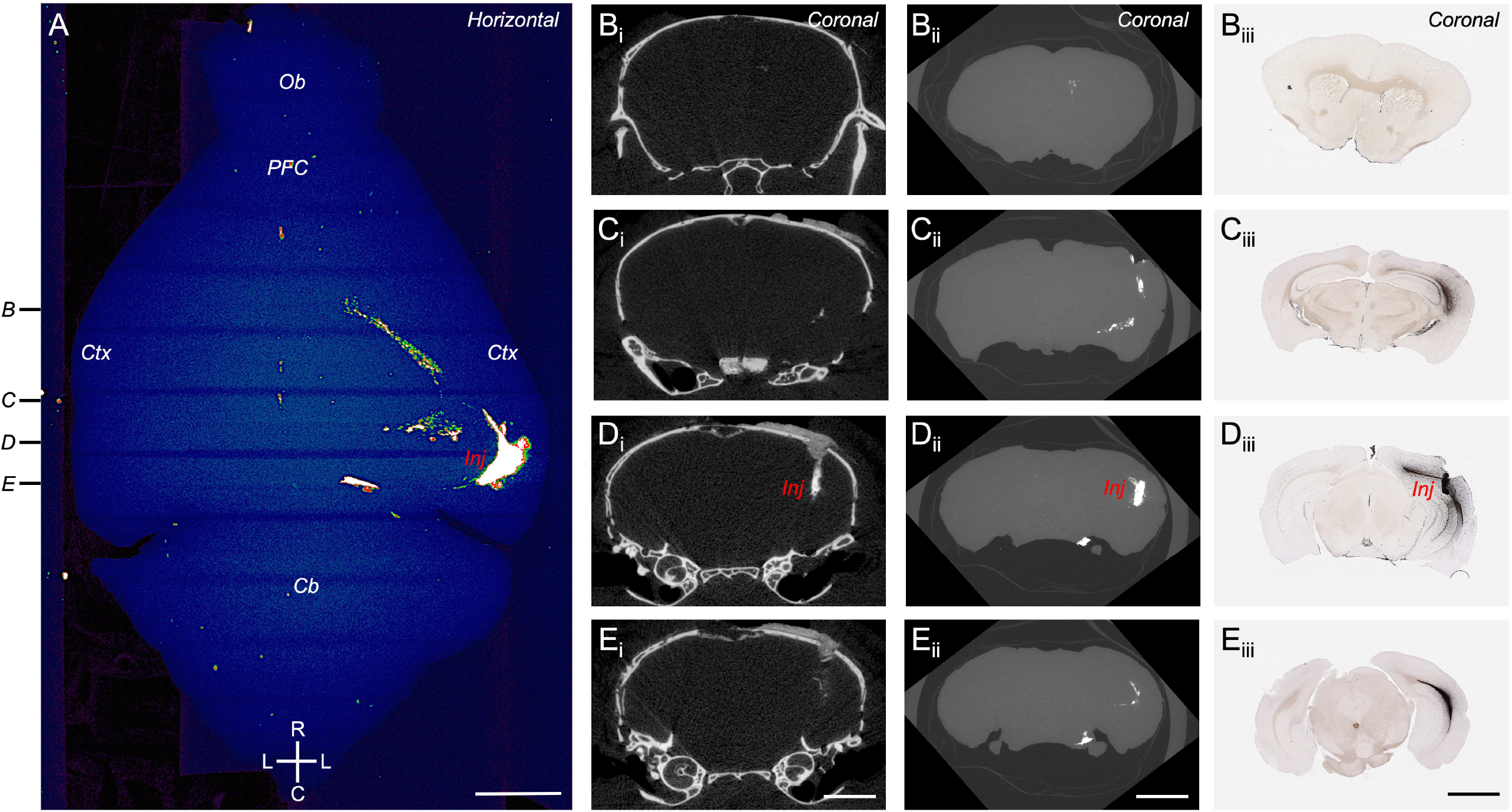
Comparison of *in vivo* and *ex vivo* micro-CT images with histologically processed sections. **A.** 3D-reconstructed micro-CT dorsal horizontal view of the mouse brain showing the injection site (*Inj*) and subsequent labeling distribution across the brain. **B-E**. Coronal cross-sectional images from *in vivo* (i), *ex vivo*, (ii), and histological (iii) specimens. See also Supplementary Video 2. Approximate rostrocaudal cross-sectional levels (*B-E*) indicated on the left of panel *A*. Scale bars: 2 mm. Orientation: C, caudal; L, lateral; R, rostral. Brain areas: Cb, cerebellum; Ctx, cerebral cortex; Ob, olfactory bulb; PFC, prefrontal cortex.

To test the degree to which the skull attenuated the detection of the WAHG tracer, we perfused and dissected the brains, and then imaged them using both the *in vivo* and *ex vivo* scanners (Figs. 2ii, 3; Suppl. Fig. 1, Suppl. Videos 2,3). We were able to identify several additional sources of labeling that were not detected in the encapsulated brains (Figs. 1-3, Suppl. Video 2,3). In addition, since the *in vivo* scanner could accommodate several brains simultaneously, we assessed whether it could be used to rapidly compare labeling in multiple brain specimens (Fig. 3, Suppl. Video 3). In this regard, we found that four brains could be rapidly and simultaneously imaged with high resolution; each of these brains showed distinct, yet related, patterns of retrograde labeling following varied hippocampal injections that could be directly compared in the same scan (Fig. 3, Suppl. Video 3).

**Figure 3.**
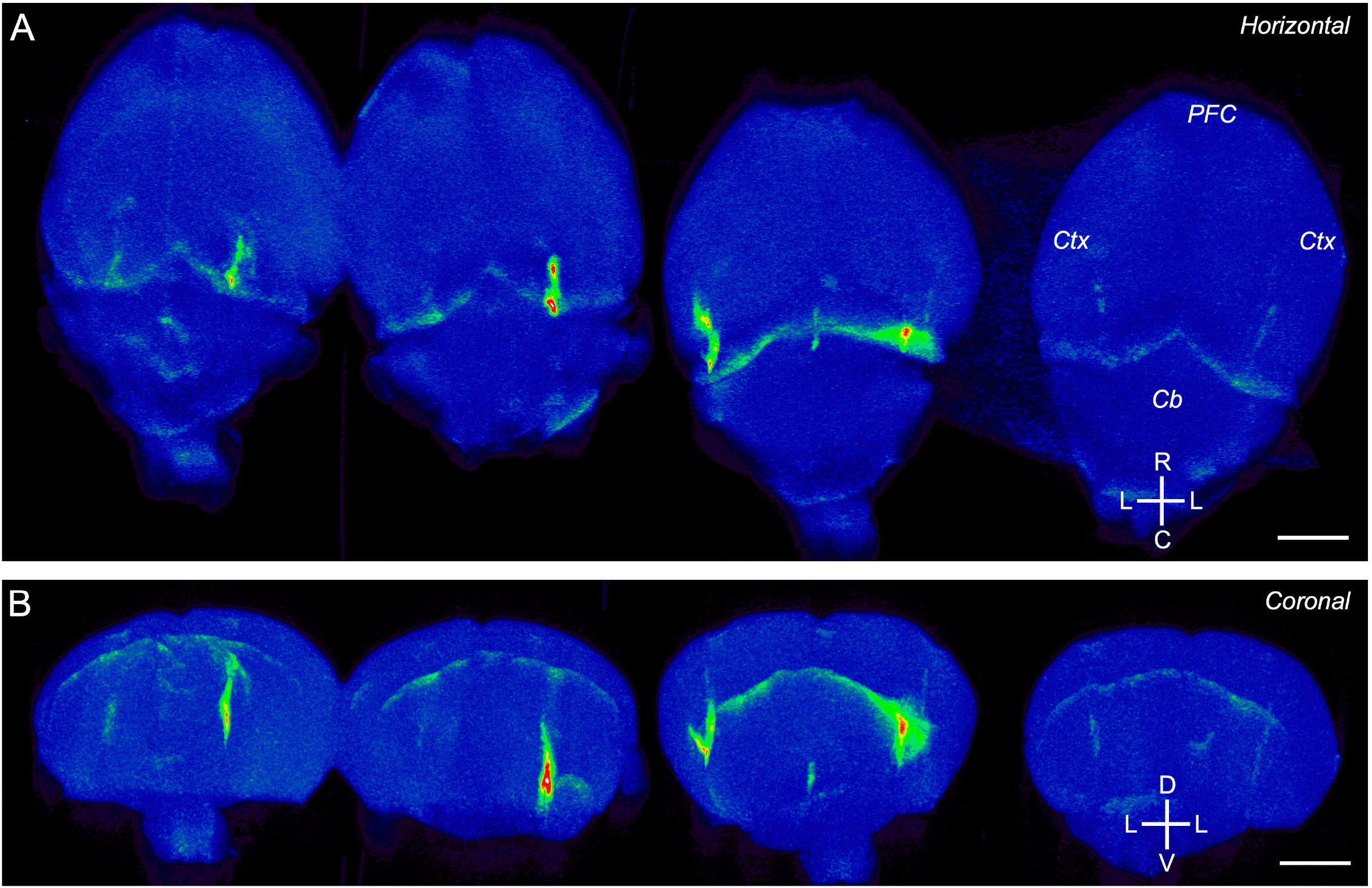
Simultaneous micro-CT scanning of injected brain specimens. **A.** 3D-reconstructed micro-CT dorsal horizontal view of the mouse brain showing the injection site (*Inj*) and subsequent labeling distribution in four different mouse brains, scanned simultaneously. **B**. The same brains in *A*, viewed from a caudal horizontal orientation. Scale bars: 2 mm. Orientation: C, caudal; D, dorsal; L, lateral; R, rostral; V, ventral. Brain areas: Cb, cerebellum; Ctx, cerebral cortex; PFC, prefrontal cortex.

Finally, we wished to compare whether the acquired micro-CT images corresponded to those obtained following standard histological processing for WAHG [28]. Therefore, after the final set of micro-CT images were obtained, we processed the sectioned tissue using silver intensification for the gold-labeled particles (Fig. 2iii). We observed extensive labeling in the ipsilateral hemisphere, with lighter labeling observed in the contralateral hemisphere (Fig. 2iii). The most intense labeling was concentrated near the injection site and in the surrounding ipsilateral hippocampal and cortical regions, which corresponded to the location of high signal detection in the micro-CT images from both encapsulated and dissected brains (Fig. 2i-iii). Histological processing also revealed a broader swath of WAHG labeling that was not visible from the micro-CT images, revealing an upper-limit on signal detectability with our current imaging equipment (Fig. 2).

## DISCUSSION

Here, we described a novel ‘neurotomographic’ imaging approach that employs micro-CT detection of a gold-labelled tracer (WAHG) for mesoscale mapping of interconnected neural structures. Our data demonstrate the potential of this approach for imaging the 3D location of injection sites and subsequent labelling in both encapsulated and dissected brains. We suggest that our novel approach can be applied to many circuit questions in systems neuroscience, such as for identifying connected networks in multi-site electrophysiological recordings and for the rapid assessment of mesoscale connectivity in several brain specimens simultaneously. In addition, this approach could be applied to the longitudinal monitoring of connectivity with age and for use in animals with much larger brains that are not as amenable to imaging with current optical clearing approaches. Finally, this micro-CT based imaging approach enables the validation of injection site placements in long-term behavioural and physiological studies, which are typically validated histologically after the conclusion of the study. As such, this method can identify subjects with either unsuccessful or misplaced injections, thus enabling an initial culling of these subjects from these long-term studies and the streamlining of such labor intensive experiments [32, 33].

Compared to currently used imaging approaches, micro-computed tomography (micro-CT) is a non-destructive imaging modality that is capable of revealing aspects of neural architecture *in vivo*. When paired with tracers or compounds that are coupled with X-ray contrast agents, it enables visualization of neural circuits in intact tissue volumes [19-22]. Since it is non-destructive, it can enable experiments that involve repeated analysis and multi-modalities. Moreover, since chemical processing steps are not required to visualize the contrast agents, the imaging can be carried out immediately, rapidly, and repeatedly.

Our novel neurotomographic approach presented here demonstrates that micro-CT coupled with WAHG tracers enables visualization of major mesoscale neural labelling patterns without physical sectioning. From our study, we propose that this approach can be refined further to improve its resolution, detectability, and cell-type specificity. First, micro-CT equipment can potentially be further optimized for these types of neural tracing studies [34, 35]. Secondly, multiple tracers can be coupled to different high-Z elements, e.g. iodine, gold, bismuth, etc., which can enable double labelling approaches [36]. These different high Z-element tracers can be resolved independently using either dual (multi)-energy X-ray excitation and/or photon counting X-ray detectors [24, 25]. Finally, virally engineered tracers can be constructed to express proteins that concentrate high Z-elements in neuronal cell bodies and their projections. Two possibilities include metallothionein (MT) and the sodium iodide symporter (NIS), which are also amenable for SPECT studies [37, 38]. Such virally encoded high Z-tracers could be engineered for cell-type specific expression, e.g. cre-lox recombination with relevant transgenic strains [39].

Overall, the novel neurotomographic approach presented here utilizes the unexplored potential of micro-CT for exploring the nervous system. As such, the approach has strong potential to complement the existing suite of approaches for investigating the structural organization of the nervous system. We propose that this novel methodology can help assay the role of convergent neural circuits and enable targeted *in vivo* manipulations of these pathways, which should eventually lead to an understanding of neural computations occurring across whole-brain networks.

## ACKNOWLEDGEMENTS

We thank Ms. Sherry Ring for assistance with histology and Ms. Aneta Staszkiewicz for microscopy assistance.

