## Supplementary figures and images for "A Neurotomographic Approach for Mesoscale Mapping of Neural Circuits"

### Supplemental Figure 1

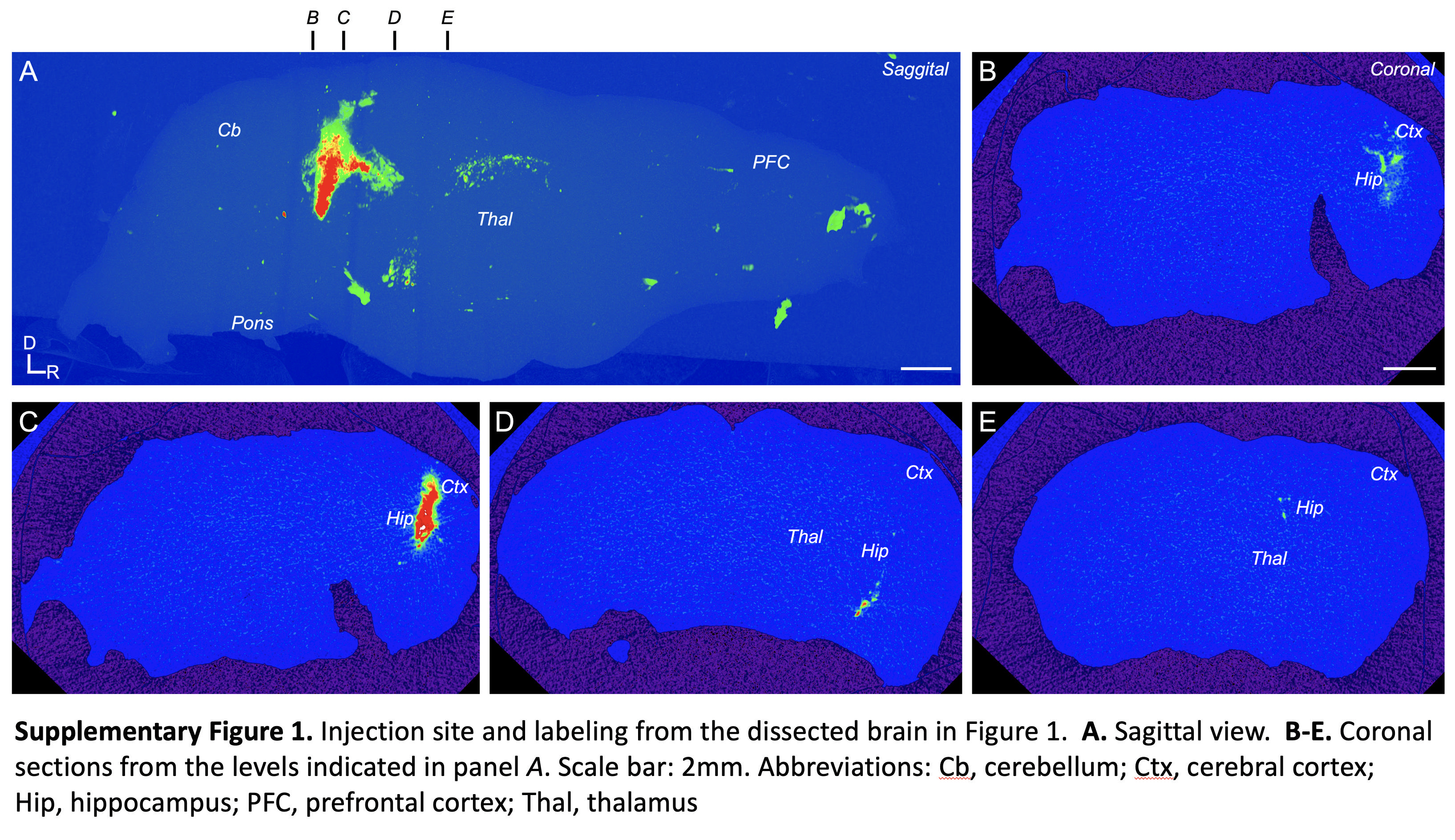
